# Cell-mechanical parameter estimation from 1D cell trajectories using simulation-based inference

**DOI:** 10.1101/2024.09.06.611766

**Authors:** Johannes C. J. Heyn, Miguel Atienza Juanatey, Martin Falcke, Joachim O. Rädler

## Abstract

Trajectories of motile cells represent a rich source of data that provide insights into the mechanisms of cell migration via mathematical modeling and statistical analysis. However, mechanistic models require cell type dependent parameter estimation, which in case of computational simulation is technically challenging due to the nonlinear and inherently stochastic nature of the models. Here, we employ simulation-based inference (SBI) to estimate cell specific model parameters from cell trajectories based on Bayesian inference. Using automated time-lapse image acquisition and image recognition large sets of 1D single cell trajectories are recorded from cells migrating on microfabricated lanes. A deep neural density estimator is trained via simulated trajectories generated from a previously published mechanical model of cell migration. The trained neural network in turn is used to infer the probability distribution of a limited number of model parameters that correspond to the experimental trajectories. Our results demonstrate the efficacy of SBI in discerning properties specific to non-cancerous breast epithelial cell line MCF-10A and cancerous breast epithelial cell line MDA-MB-231. Moreover, SBI is capable of unveiling the impact of inhibitors Latrunculin A and Y-27632 on the relevant elements in the model without prior knowledge of the effect of inhibitors. The proposed approach of SBI based data analysis combined with a standardized migration platform opens new avenues for the installation of cell motility libraries, including cytoskeleton drug efficacies,and may play a role in the evaluation of refined models.

**Subject Areas:** Biological Physics / Interdisciplinary Physics

## Introduction

Cell migration on one-dimensional (1D) microlanes has become a well established cell motility assay, offering comparability, reproducibility and high-throughput automation (1–7). The reduction of cell movies to low-dimensional trajectories enables the characterization of cell populations in terms of statistical measures including cell-cell variability both within and between diverse populations. A remarkable feature in the analysis of cell dynamics is the fact that morphodynamics exhibit both cell-specific as well as universal behaviors. In early work the mean velocity of cells migrating on flat substrates was frequently studied as a cell-specific property. The average cell speed is understood to be a quantity dependent on cell line, individual cell state as well as varying external conditions. In contrast, the dependence of the speed of cells migrating on substrates as a function of increasing adhesiveness exhibits a recurrent biphasic adhesion-velocity relation that holds for many cell types (8–12). Moreover, the persistent random walk model proved to generally reproduce the diffusive nature of cell walks over large time scales. In the case of 1D migration on micropatterned microlanes, a persistent random walk analysis has led to the discovery of another universal relation, the so called universal coupling of cell speed and persistence (UCSP) (2,13). A long history of cell migration models aimed to explain the underlying cause of universal features in cellular morphodynamics. In particular, detailed biomechanical models of migration in one dimension have been developed to elucidate the features observed in cell trajectories (4,12–17). These models exhibit a rich spectrum of behavior, including multiple cell states with distinct dynamic features, specifically states with oscillations of the rear end versus steady state motion. The characteristics of states as well as the noise driven transition statistics between states are cell type specific. In order to validate mechanistic models, comparison to experimental data is requested and requires an optimal choice of parameters in the theoretical models. The majority of models have been validated using a limited number of cell lines and parameter sets. The complexity of biomechanical models is demanding and the effect of parameter changes is not always intuitive. As a consequence parameter optimization is both mathematically and conceptually challenging and the high-dimensionality of the problem makes rigorous Bayesian inference computationally infeasible. Often, researchers are left to explore model parameters based on intuition or trial-and-error in a laborious and non-systematic way. A systematic and scalable approach to infer parameters from large cell motility datasets would allow for data-driven discovery and extraction of information about the underlying networks regulating cell motility.

In recent years, machine learning (ML) approaches have emerged as a powerful tool to analyze cell phenotype, including cell morphology and dynamics. Neural networks proved useful for automated image segmentation and retrieval of cell shape from phase contrast or fluorescence image raw data (18,19). Early cell shape analysis approaches used classic Fourier analysis for the classification of cell shape dynamics (20). In recent years deep learning methods led to robust cell type classification schemes and recognition of disease related morphometry (21–25). Furthermore, AI based approaches enable data-driven discovery from large biological data sets of cell shapes under defined conditions(26). Using real-time assessment of cell shapes in standardized platforms a novel type of cytologic analysis emerged, an approach now commercialized under the name “morpholomics” (27–29). However, there are few AI-based approaches that include the dynamic features of cell shapes. In dynamic analysis the reduction of cell motion to one dimension helps to reduce the complexity of cell morphodynamics.

In the latter case mechanistic models of dynamical behavior exist and deep learning approaches offer the unique opportunity that neural density estimators can be trained on simulated data. The trained network in turn is capable of estimating best model parameters to fit experimental data and hence offers new avenues for parameter optimization in complex models using Bayesian inference (30). The approach named simulation-based inference (SBI), has gained widespread acceptance as a systematic tool for parameter optimization in mechanistic models (31–34). A particularly inspiring example is the Hodgkin-Huxley model reproducing neural spikes, in which case SBI is used to estimate model parameters to capture specific experimental spike recordings (31). SBI combines elements of simulation modeling and statistical inference to analyze data and make inferences about underlying processes. In order to train SBI, parameter sets are sampled from the prior, i.e. a set of candidate values, to simulate data using the model. Next, a deep density estimation neural network is trained to infer the parameters underlying the simulated data. Finally, the trained density estimation network is applied to experimental data to infer its parameter distribution. SBI is particularly useful in situations where: the underlying system is complex, with intricate interactions and dependencies that are difficult to model analytically; traditional likelihood-based methods may not be feasible due to intractable likelihood functions or computational limitations; the data exhibit heterogeneity or non-standard patterns that cannot be adequately captured by standard statistical models; there exists prior knowledge or mechanistic understanding of the system, which can be incorporated into the simulation process. Despite its general applicability and statistical power, the application of SBI to derive migratory phenotypes with specific model parameters has been limited. In this context we recently presented a mechanistic model based on the concept of competing protrusions and noisy clutch. Within this model universal relations, such as the adhesion-velocity relation and the UCSP which is caused by multistability, emerge as embedded features of the nonlinear dynamics (12,13). However, a model based characterization of migration dynamics across multiple cell lines, each distinguished by specific sets of parameters, has yet to be done.

Here, for the first time, we employ SBI to infer parameter sets from 1D trajectories of migrating cells within the framework of an established mechanical migration model (13). We train a neural network using simulated trajectories and infer parameters from high-throughput datasets containing hundreds of experimentally obtained trajectories. Using an algorithm based on work by Papamakarios and Murray, Lueckmann et al., Greenberg et al. and Deistler et al. (35–38), we first validate our approach on simulated data before we systematically optimize the parameter set matching the migratory behavior of two human epithelial breast cell lines. Specifically, we obtain cell-line-specific posterior estimators for cell length, actin polymerization rate and the integrin related parametrization of the clutch, i.e. the on-rate, slip velocity and maximum friction coefficient. We demonstrate that the estimated parameters significantly vary for cell lines MDA-MB-231 and MCF-10A. SBI is also found to unveil without prior knowledge that the cytoskeletal inhibitors Latrunculin A and Y-27632 both exclusively affect actin polymerization. Our work showcases the potential of SBI to characterize migrating cells in a fully automated fashion and to explore the compliance of refined biophysical models.

## Results

### High-throughput imaging of 1D cell migration yields large amounts of data to quantitatively study migratory behavior

We study epithelial cells on a micro-patterned substrate consisting of well-defined adhesive Fibronectin (FN) lanes separated by nonadhesive regions, see **Fig 1(a)**. After the cells adhere to the FN lanes, we monitor cell migration via automated scanning time-lapse acquisition over 48h (see Methods). For statistical analysis a high number of cell trajectories of substantial length is required. From previous work it is known that migratory behavior shows substantial heterogeneity even within the same cell population and under controlled conditions. Furthermore the analysis of rare events such as migratory state transitions, require a long total observation time (13). We met this requirement by building a data pipeline to automatically extract cell trajectories from time-lapse videos imaging a few thousand cells per experiment. The pipeline takes a set of time-lapse images and automatically outputs a dataset with several thousand triple trajectories consisting of the front, back and nucleus position of single cells for each experiment. For our analysis we consider only single cell trajectories with a minimal length of 24h. A total of 47,000hours of tracks is automatically processed. We collected about 2,000 single cell trajectories of 24h length for both the breast epithelial cell lines MDA-MB-231 and MCF-10A. In **Fig 1(b)** we show typical trajectories for each cell line. The spectrum of migratory phenotypes is broad, with trajectories moving at a range of different speeds or not moving at all, as well as protrusions oscillating frequently in length or staying stationary. The data set was complemented by studies of cells treated with inhibitors latrunculin A, which inhibits F-actin polymerisation, and with Y-27632, which inhibits Rho-associated protein kinase (ROCK) signaling (39,40). We chose the concentrations for the treatments to be high enough to affect the migratory behavior but low enough to not stall migration altogether (see Methods). For this study the large-scale acquisition of 1D trajectories provides the biological information about the migratory phenotype of cells. However, in order to retrieve an understanding of phenotypic characteristics a mechanistic model is essential.

**Fig 1.**
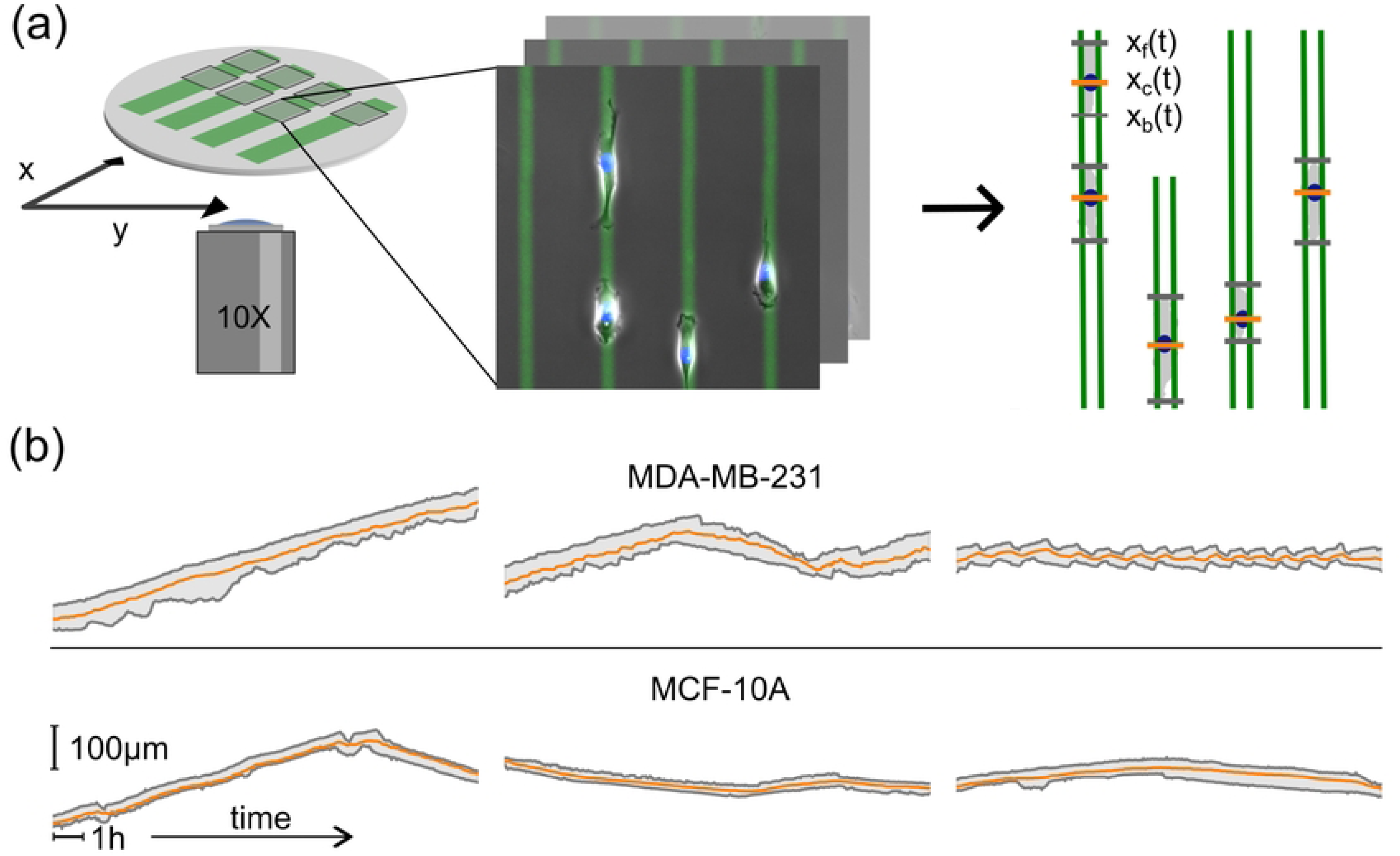
High-throughput time-lapse imaging of single cells migrating on 1D micropatterned lanes. **(a)** Migration dynamics of single cells on one-dimensional Fibronectin (FN) lanes is recorded by scanning time-lapse measurements. FN lanes are fluorescently labeled, depicted here in green. At each point in time a bright field (BF) image showing the cell contour, and a DAPI fluorescence image indicating the position of the nuclei are captured. The FN lanes are automatically detected, the nuclei tracked and the cells’ contours segmented. **(b)** Single cell trajectories, shown as position of cell front, back and nucleus plotted against time, reveal a broad spectrum of cell-type specific features. The first row depicts typical trajectories for MDA-MB-231 cells, the second row for MCF-10A cells. Horizontal scale bar represents 1h, vertical scale bar 100um.

### Biophysical model of mesenchymal cell motility

In previous work we introduced a biophysical model that reproduced all the observed universal migratory hallmarks, including multistability of migratory states, the universal correlation between speed and persistence (UCSP) and the biphasic adhesion-velocity relation (13). We use this model as a candidate model for SBI with the goal to characterize the cell-specific migration dynamics of MDA-MB-231 and MCF-10A cells. The model describes a cell that migrates along a lane by a one-dimensional mechanical equivalent model consisting of a nucleus flanked by a lamellipodium of length L on each side (Fig 2). Lamellipodia have the resting length L_0_ in the force-free state and are coupled to the nucleus by springs with an elastic modulus E. The elastic forces mediate competition between the protrusions. Protrusion forces F_f_ and F_b_ arise from the extension of the F-actin network at rate V_e_ which pushes against the cell’s front and back edges by polymerization of filament tips near the cell membrane. This polymerization force also drives retrograde flow v_r_ against the friction force F_fric_ = ⍰*v_r_. Friction force is caused by the binding and unbinding of the actin retrograde flow to structures adhered to the substrate which are stationary in the lab frame of reference with rates k_on_ and k_off_(v_r_), respectively (41,42). The dissociation rate depends on the velocity of the retrograde actin flow v_r_ in a non-linear fashion. The dynamics can be characterized by k_on_, k_off_, the maximum friction coefficient ⍰_max_ and the characteristic retrograde flow velocity v_slip_, see SI. Noise ε adds the stochastic behavior observed in experiments. The cell’s edges and nucleus experience a drag force that depends linearly on the cell’s velocity v with the drag coefficient ⍰. In our previous work, the drag coefficient ⍰ was related to the fibronectin density B by Hill-type equations with the maximum ⍰_max_, see SI. As the fibronectin density is constant in the present study we simplify the model and introduce a constant B. We keep the ratio b of the drag at the cell’s nucleus versus that of the edges constant, too. Lastly, we add an external noise term ε_external_ to the model to account for random motion on short time scales that might, among other things, be caused by limitations of the experimental determination of the cell’s location (see Methods). Computational simulations based on this model reproduce simulated trajectories that resembled observed data as indicated in Fig 3. In order to determine the parameter sets that show best agreement with data, we employ simulation-based inference as explained in the next section.

**Fig 2.**
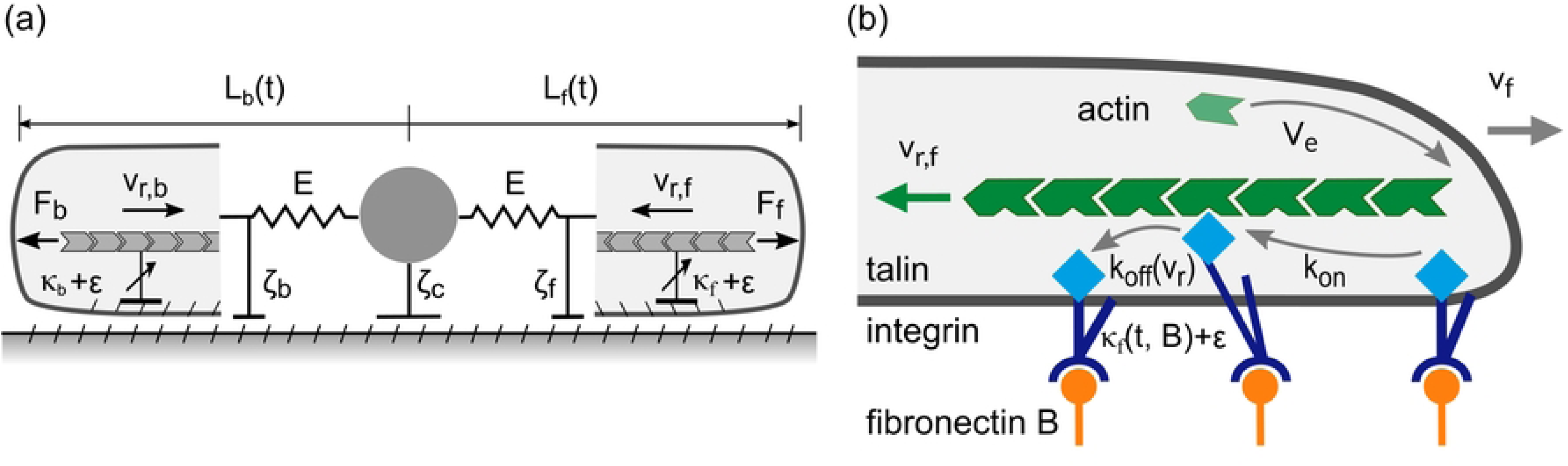
Biophysical model of cell migration in one dimension. (a) Cartoon of the mechanical protrusion competition model. The cell is defined by three marks: back, nucleus and front. Front and back are coupled to the nucleus by an effective elastic spring and coupled to the ground by a non-linear molecular clutch. Redrawn from (13). (b) Cartoon of the molecular clutch. Actin polymerisation at the edge of the cell creates a retrograde flow v_r_. Talin-integrin mediated coupling between the actin network and the fibronectin substrate results in an effective friction force. Friction slows down the retrograde flow v_r_ and pushes the membrane outwards.

**Fig 3.**
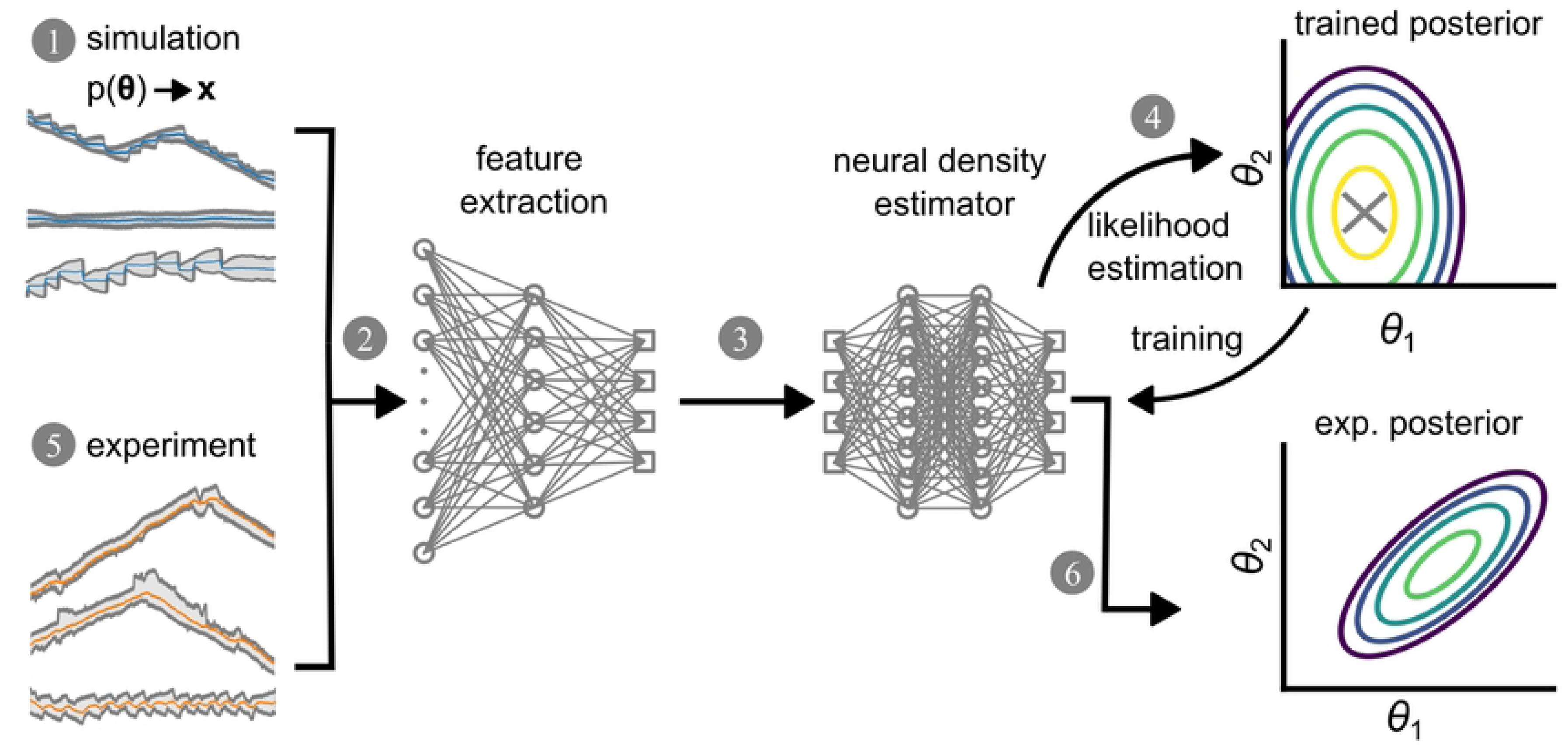
Simulation-based inference (SBI) of model parameters. A schematic representation of the SBI workflow. A neural network is trained with simulated data and subsequently experimental data are analyzed using the pretrained network. (1) A set of parameters {**θ**} is randomly sampled from the prior p(**θ**) and used to simulate trajectories **x**. (2) The trajectory is downsampled into a low-dimensional feature space by a convolutional neural network (CNN). (3) The downsampled trajectory is fed into the neural density estimator (NDE) which outputs the posterior density. The log-likelihood of the NDE at the true point (X) is used as a loss function to update the NDE. The trained NDE has a maximum likelihood at the true parameter point as marked by ‘X’. The trained SBI is then used to estimate parameters from measured data. (5) The experimental trajectories are downsampled into a low-dimensional feature space by the same CNN as in step (2) and then fed into the previously trained NDE. (6) The resulting posterior represents a cell-specific parameter estimation describing interpretable properties of the cell as defined by the biophysical model.

### Simulation-based inference connects experimental trajectories to biophysical parameter distributions

We would like to establish the relation between cell trajectories as shown in Fig. 1b and the biophysical model (Fig 2). As shown in Fig 3, our goal is to present an unbiased approach to estimate those parameters that best match the data. We will show that SBI allows for inference of the set of cytoskeletal parameters **θ** which are most likely to generate a trajectory **x**(t) using Bayes’ theorem. Bayes’ theorem states that the desired probability distribution of parameters is the posterior distribution p(**θ**|**x**) given the prior parameter distribution p(**θ**) (see Table 1), the likelihood p(**x**|**θ**) and the evidence p(**x**):

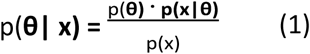

**Table 1.** Prior p(θ). The 10 parameters and two noise amplitudes entering our biophysical model. 5 parameters are variable and the target of our inference procedure; one of the noise amplitudes is latent; all other parameters are kept constant.

| Parameter name | Description | Lower bound | Upper bound | Status | Units |
| --- | --- | --- | --- | --- | --- |
| $L_0$ | Resting protrusion length | 1 | 40 | Variable | $\mu\text{m}$ |
| $V_e^0$ | Actin network extension rate | 1e-3 | 8e-2 | Variable | $\mu\text{m s}^{-1}$ |
| $k_{on}$ | On rate for dynamic integrin signaling | 1e-5 | 1e-3 | Variable | $\text{s}^{-1}$ |
| $v_{slip}$ | Critical retrograde flow velocity | 5e-3 | 4e-2 | Variable | $\mu\text{m s}^{-1}$ |
| $\zeta_{max}$ | Maximum friction coefficient for integrin signaling | 1 | 70 | Variable | $\text{nN s } \mu\text{m}^{-2}$ |
| $\eta_{\max}$ | Maximum drag coefficient | 1.4 | 1.4 | Fixed | nN s $\mu\text{m}^{-2}$ |
| E | Effective E-modulus | 3e-3 | 3e-3 | Fixed | nN $\mu\text{m}^{-2}$ |
| $k_{\text{off}}$ | Off rate for dynamic integrin signaling | 0.5 | 0.5 | Fixed | s-1 |
| $b = \eta_c/\eta_f$ | Contribution of the cell body to the cell drag compared to the protrusions | 3 | 3 | Fixed | |
| B | Fibronectin density | 30 | 30 | Fixed | ng cm-2 |
| $\epsilon$ | Noise in η-dynamics | 1 | 1 | Fixed | |
| $\epsilon_{\text{external}}$ | Noise in x position | 0.5 | 2 | Latent | $\mu\text{m}$ |

Computing the posterior distribution p(**θ**|**x**) therefore implies computing both the likelihood p(**x|θ**) and the evidence p(**x**), which is computationally infeasible for a high-dimensional parameter space. We therefore use neural density estimation (NDE) to learn the posterior distribution p(**θ**|**x**) directly without computing the likelihood p(**x**|**θ**). The algorithm itself is based on work by Papamakarios and Murray, Lueckmann et al., Greenberg et al. and Deistler et al. (35–38). Specifically we deploy the toolkit “sbi”, a PyTorch-based package developed by Tejero-Cantero et al. (43). The following procedure to implement SBI is based on the work by Goncalves et al. (31).

**Fig 3** shows the 6-step workflow of simulation-based inference using NDE. First, the algorithm randomly samples a set of points from the prior parameter space {**θ**} to simulate a set of trajectories **x** using our biomechanical model. Simulated trajectories vary in their appearance even if they are simulated using the same parameter set because of the stochastic nature of the model. Each trajectory consists of 3×721 = 2,163 data points, representing the position of the front, back and nucleus of a cell for 721 time points which signify a temporal resolution of 2min in 24h of simulated time. Second, an embedding in the form of a convolutional neural network (CNN) compresses each trajectory and extracts summary statistics, also called “features”. Third, these features are fed into a neural density estimator based on neural spline flows, a form of normalizing flows, to calculate the posterior directly. Fourth, both networks, i.e. CNN and NDE are trained on simulated trajectories with known parameters by adjusting their weights to maximize the log-likelihood of true parameters (see Methods). Fifth, the trained SBI can then be applied to empirical data from *in vitro* experiments. The experimental trajectories are treated the same way as the synthetic data, i.e. they are fed into the trained CNN and NDE, resulting in a posterior distribution. This posterior distribution assigns a likelihood to all parameter values depending on the data and the prior. Hence, SBI can estimate the optimal set of model parameters that characterizes an experimental trajectory.

Table 1 shows the 10 parameters and 2 noise amplitudes that enter our simulation as defined by the mechanical model. However, inferring the full set of parameters we encounter loss of identifiability To demonstrate the problem we discuss the results of SBI with 10 free parameters, which exhibits correlations and lacks precise inference (see SI). In order to proceed we reduce the complexity of the neural posterior estimation by rescaling our model to five most influential parameters, (for details see SI). We find that our model’s dynamics is fully described by the reduced set of the following 5 parameters: {resting protrusion length: L_0_, actin network extension rate: V_e_^0^, on-rate for dynamic integrin signaling: k_on_, maximum friction coefficient for integrin signaling: ⍰_max_, critical retrograde flow velocity: v_slip_}. As shown in the next section, the reduced set of parameters is inferred reliably without loss of identifiability.

### Validation of SBI using simulated data

We start our analysis by training an NDE using 1,000,000 simulated trajectories with known parameters to infer the 5 parameters of choice. For details we refer to the methods section. Next, we show that our posterior is well calibrated, i.e. neither underconfident nor overconfident, by performing a simulation-based calibration (SBC), see S2 Fig. The details of the procedure are described in the SI. We then validate the performance of our NDE by testing its predictive power using simulated data and by making sure that it is unbiased. To this end we generate a test trajectory from randomly chosen but known parameters and subject the data to SBI. **Fig 4** shows the simulated trajectory **x**(t) and the resulting posterior p(**θ**|**x**) as well as the true parameter **θ_true_** set used to simulate the trajectory. The parameter set for the simulated trajectory is randomly sampled from a uniform prior distribution. The trajectory simulates a cell that migrates along a 1D FN lane for 24h without the influence of any external forces. The simulated trajectory exhibits several changes in the direction of the cell’s movement and an oscillating length of the cell. By visual inspection we see that the inferred posterior generally peaks at the values of the true parameters (marked by a vertical orange line) which indicates an accurate and unbiased inference. Plots of the pairwise distribution p(θ_ij_|x) in the right hand corner of Fig 4 provide insights into correlations between parameters. Tilted distributions indicate a positive or negative (depending on the sign of the slope) correlation, while horizontal distributions, such as for k_on_ vs V_e_^0^, indicate the parameters to be orthogonal. Clearly, the sensitivity of the inference of parameters varies. Parameters that can be particularly accurately estimated, as can be seen by a sharp prior distribution, are the resting cell length L_0_ and the network extension rate V_e_^0^. In summary, applying a simulated trajectory with known parameters to the trained NDE correctly infers posterior distributions for 5 free parameters that comprise the wanted parameters within the accuracy of the approach.

**Fig 4.**
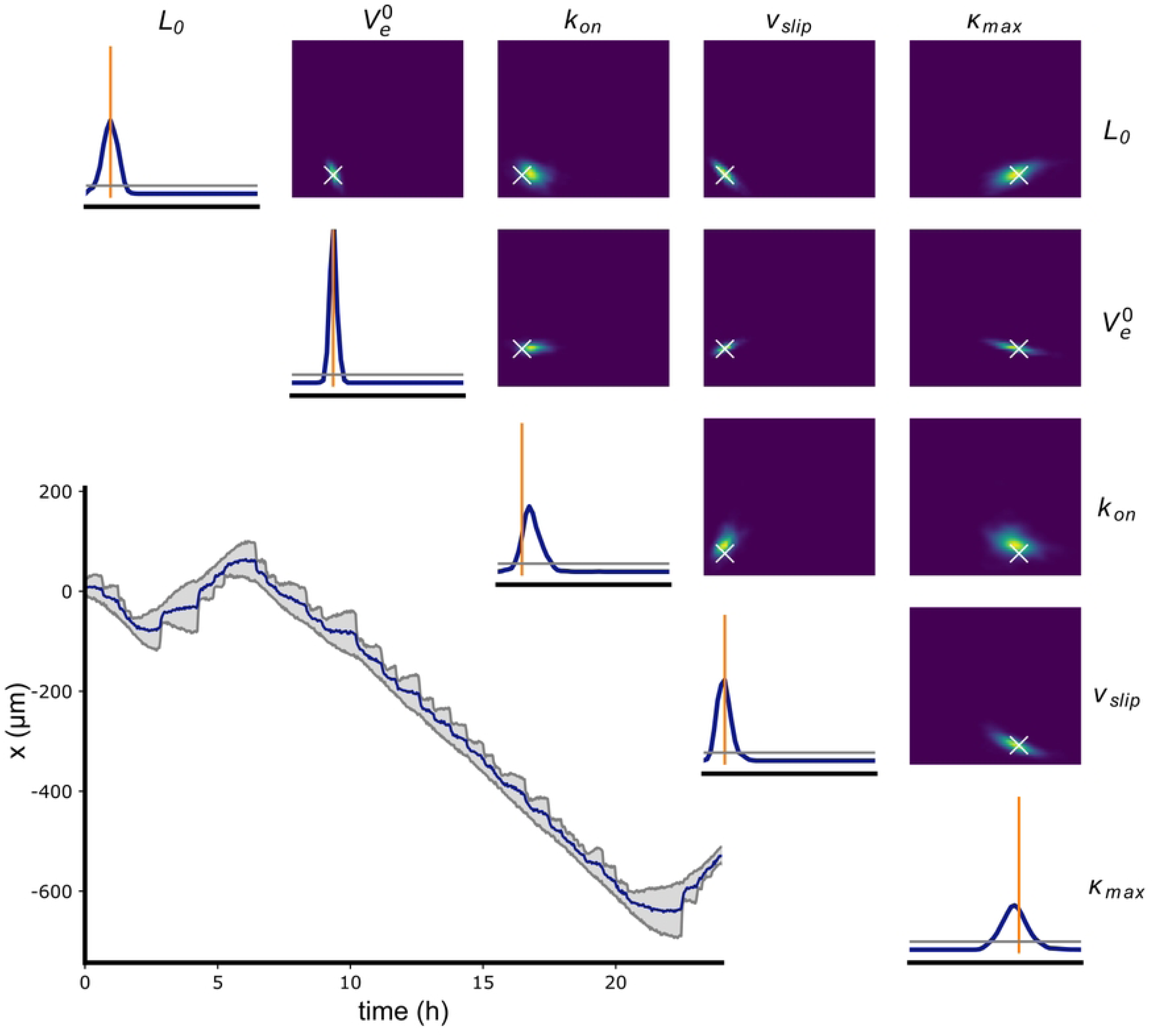
Validation of SBI applied to a simulated trajectory. A simulated trajectory **x** from parameter set **θ** and its corresponding posterior distribution. The posterior probability p(**θ**|**x**) was inferred by the trained NDE. On the right hand side the posterior distributions for each pair of parameters p(θ_ij_|x) are plotted (true parameters shown as white cross). The posterior distribution p(θ_i_|x) for each individual parameter estimation is given by the blue graph on the diagonal with the true parameter value indicated by the vertical orange line. Horizontal gray lines in the plots on the diagonal represent a uniform posterior. The boundaries of the x-axis correspond to those of the prior p(θi) (see Table 1).

### Inference of parameters from experimental trajectories

Next, we apply the trained NDE to experimental trajectories of MDA-MB-231 cells in two different states of motion as depicted in **Fig 5**. The insets of panel (a) and (b) show the trajectories, with trajectory (a) constantly moving albeit at different speeds and trajectory (b) being spread for most of the time. While the lengths of the protrusions for trajectory (a) oscillate for the first couple of hours, they stay relatively constant after 12h. Trajectory (b) on the other hand represents a cell whose protrusions keep on oscillating in length for the entirety of the observed time. The diagonals in **Figs 5(a)** and **5(b)** display the inferred posterior distributions for each individual parameter and the right hand corners display distributions for pairs of parameters. The posterior distribution of actin network extension rate V_e_^0^ is strongly peaked for both trajectories while the distribution for v_slip_ is very broad and close to that of a uniform posterior distribution (horizontal gray line). The inferred distribution of L_0_ is much broader for trajectory (a), where the protrusion length and oscillatory behavior changes over time, compared to that of trajectory (b), which oscillates permanently. Insets in the right panel of **Figs 5(a)** and **5(b)** depict simulations that were sampled from the most likely parameter set of trajectory (a) and (b), respectively. The example demonstrates that SBI is capable of inferring probability distributions of parameters for individual cell trajectories. Next, we show that parameter sets inferred for populations of different cell types result in a meaningful characterisation that discriminates distinct cell lines.

**Fig 5.**
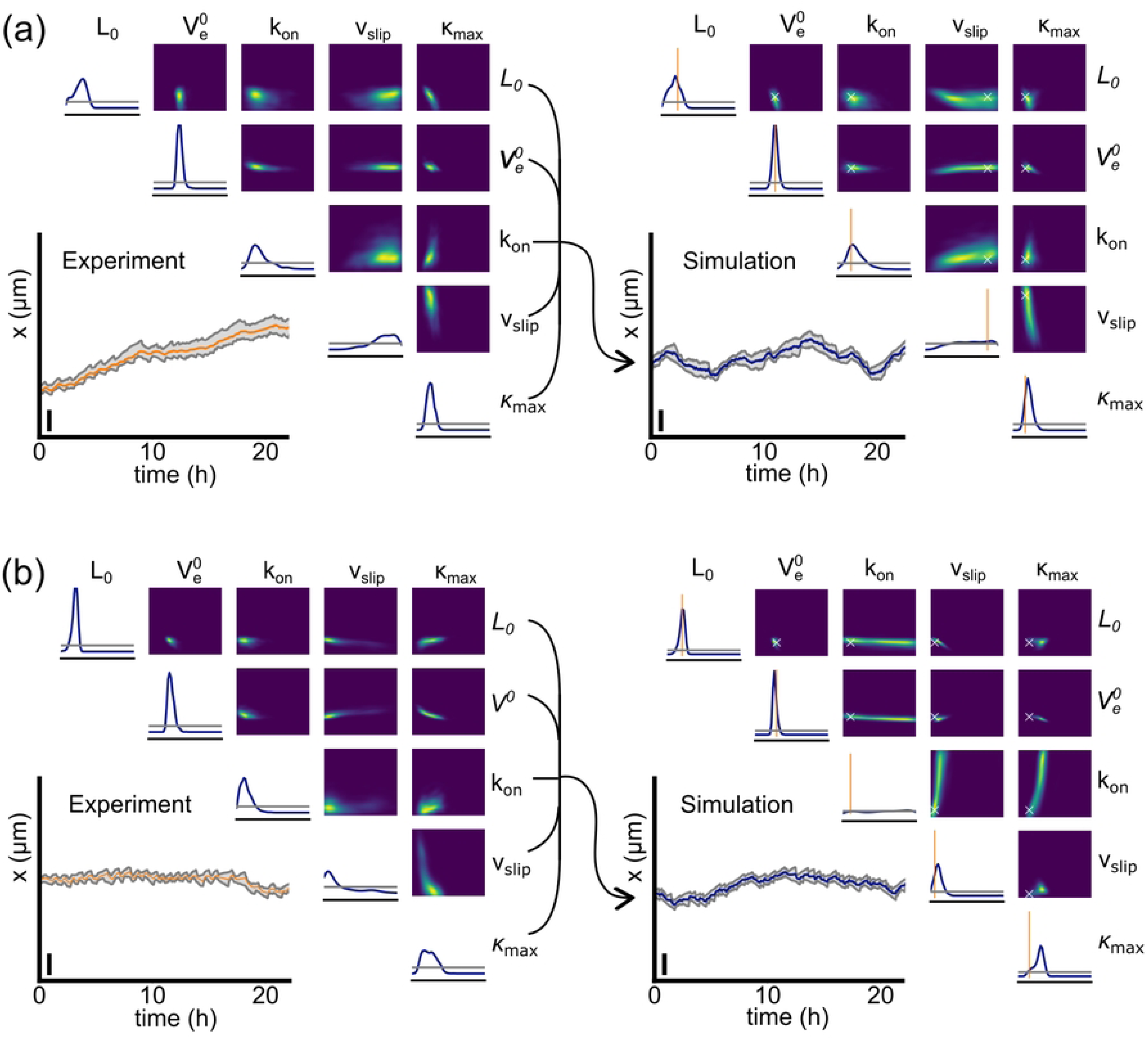
Validation of SBI on experimental data. Four different trajectories and posterior probabilities p(**θ**|**x**) as inferred by our trained NDE from experimental (left side) and simulated data (right side). The trajectories can be seen in the lower right corner. The posterior distribution for each individual parameter p(θ_i_|x) are plotted on the diagonals and in the upper right corner the posterior distribution for each pair of parameters p(θ_ij_|x) are shown. Left side in **(a)** and **(b)**: typical 24h trajectories of MDA-MB-231 cells and their estimated posterior distribution. Vertical scale bars represent 100um. Right side: the posterior distributions corresponding to each of the two experimental trajectories were used to sample parameters {**θ**}. The most likely parameter value from the experimental posterior distribution **θ_true_** was chosen to simulate a trajectory **x**, and the simulated trajectory **x** used to estimate the posterior distribution again to verify our approach. The vertical orange lines on the histograms and the crosses in the density plots show **θ_true_**.

### Inference of cell type specific properties

We use the trained NDE to characterize datasets of two different cell lines MDA-MB-231 and MCF-10A. For each population, we filtered for trajectories with a duration of 24h. Shorter durations had previously resulted in broad posterior distributions with smeared out peaks. The posterior distributions of each trajectory were used to build a 5-dimensional probability distribution of parameter values. The distributions of trajectories belonging to each cell population were combined to construct an ensemble distribution of cytoskeletal parameters for the given population, see **Fig 6**. We find that the populations of MDA-MB-231 and MCF-10A differ mainly in the distribution of the two parameters L_0_ and V_e_^0^, with the actin network extension rate being significantly higher for MDA-MB-231. Both populations express a broad, almost uniform distribution for the parameters k_on_ and v_slip_. The distribution of ⍰_max_ is similarly peaked for both cell lines, hinting towards a well conserved signaling pathway across cell lines. A comparison between 10 randomly chosen trajectories of each cell line visualizes the apparent differences in motile behavior, **Fig 6(a,b)**. While both populations exhibit both motile and spread cells, the ratio of motile cells is higher for MDA-MB-231 cells. Additionally, MDA-MB-231 cells tend to oscillate in length significantly more often than MCF-10A cells. These observed differences are explained by differences in the force-free resting length L_0_ and the actin network extension rate V_e_^0^. According to our biophysical model a shorter length and a higher actin network extension rate result in less persistent cells that are more likely to exhibit length oscillations. Hence, inference of 5 cell type specific model parameters allows for an automated and unbiased characterization of cell properties. The most distinctive cell parameters appeared to be the resting length L_0_ and the actin network extension rate V_e_^0^.

**Fig 6.**
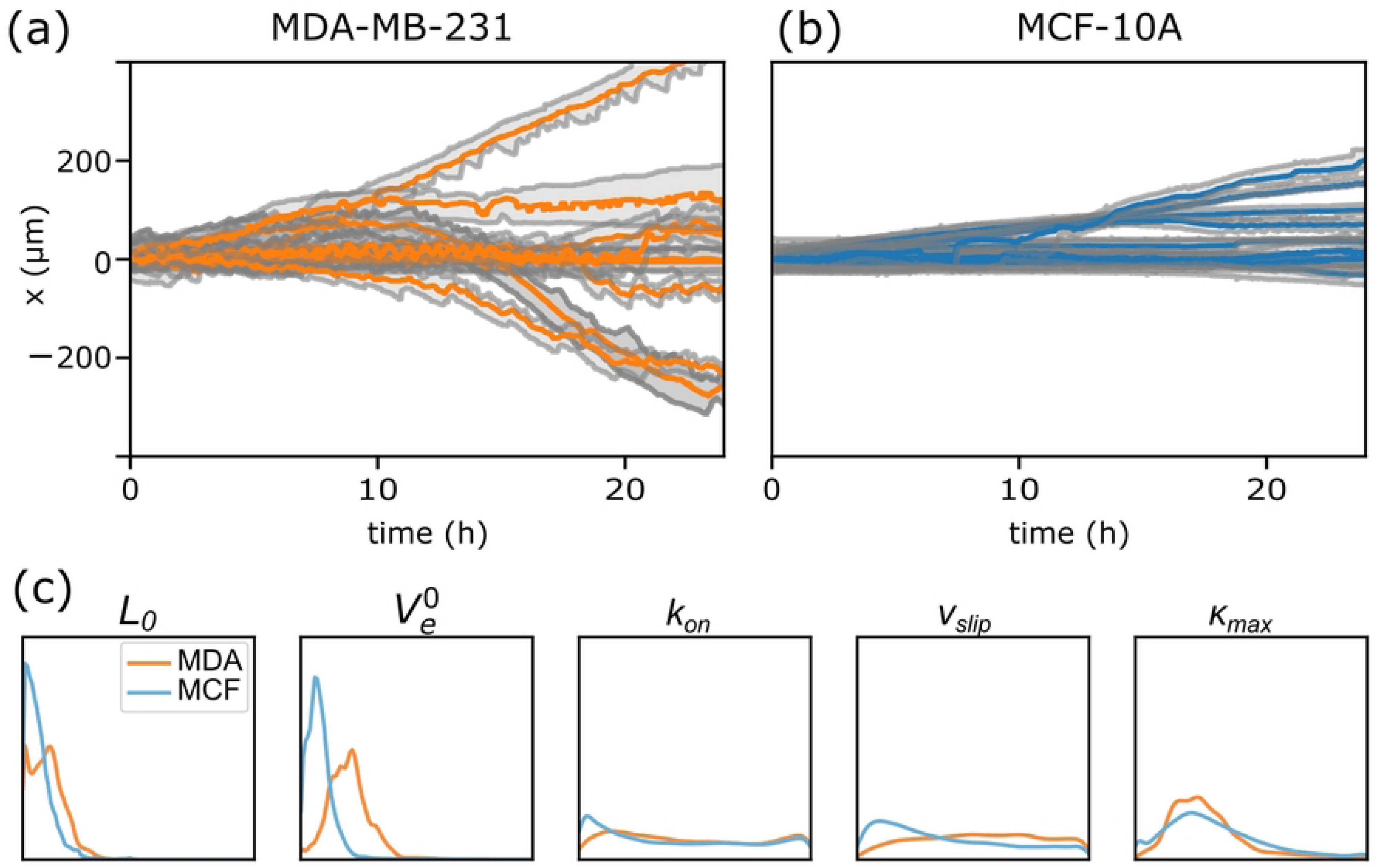
Comparative characterization of migratory phenotype for two cell lines. (**a, b)** 10 randomly chosen trajectories for MDA-MB-231 and MCF-10 cells, respectively. **(c)** Ensemble posterior distribution of estimated model parameters using SBI. For each trajectory in a given population 1000 different points were sampled in parameter space. The plots show the ensemble average of all sampled points for all trajectories of a given population (N_MDA_ = 85, N_MCF_ = 30). Cell length L_0_ and actin polymerization rate V_e_ ^0^ are the most distinctive parameters.

### Unbiased SBI analysis of the effect of inhibitors

To further test the capabilities of SBI, we subject both cell lines to cytoskeleton inhibitors. Latrunculin A inhibits the polymerisation of F-actin (39,44,45); the specific ROCK (Rho-associated protein kinase) inhibitor Y-27632 affects the Rho/ROCK pathway (46–48). We apply SBI, as described above, without implementation of prior knowledge of the inhibitor action, to the data sets. Upon treatment with Latrunculin A the inferred posterior distributions show an exclusive reduction in the rate of actin polymerization (V_e_^0^) compared to the untreated cohort both in MDA-MB-231 cells and in MCF-10A cells, **Fig 7(a)**. The inhibitor Y-27632 shows similar reduction in the polymerization rate, but additionally shifts the probability distribution of the resting protrusion length L_0_ towards larger values, **Fig 7(b)**.

**Fig 7.**
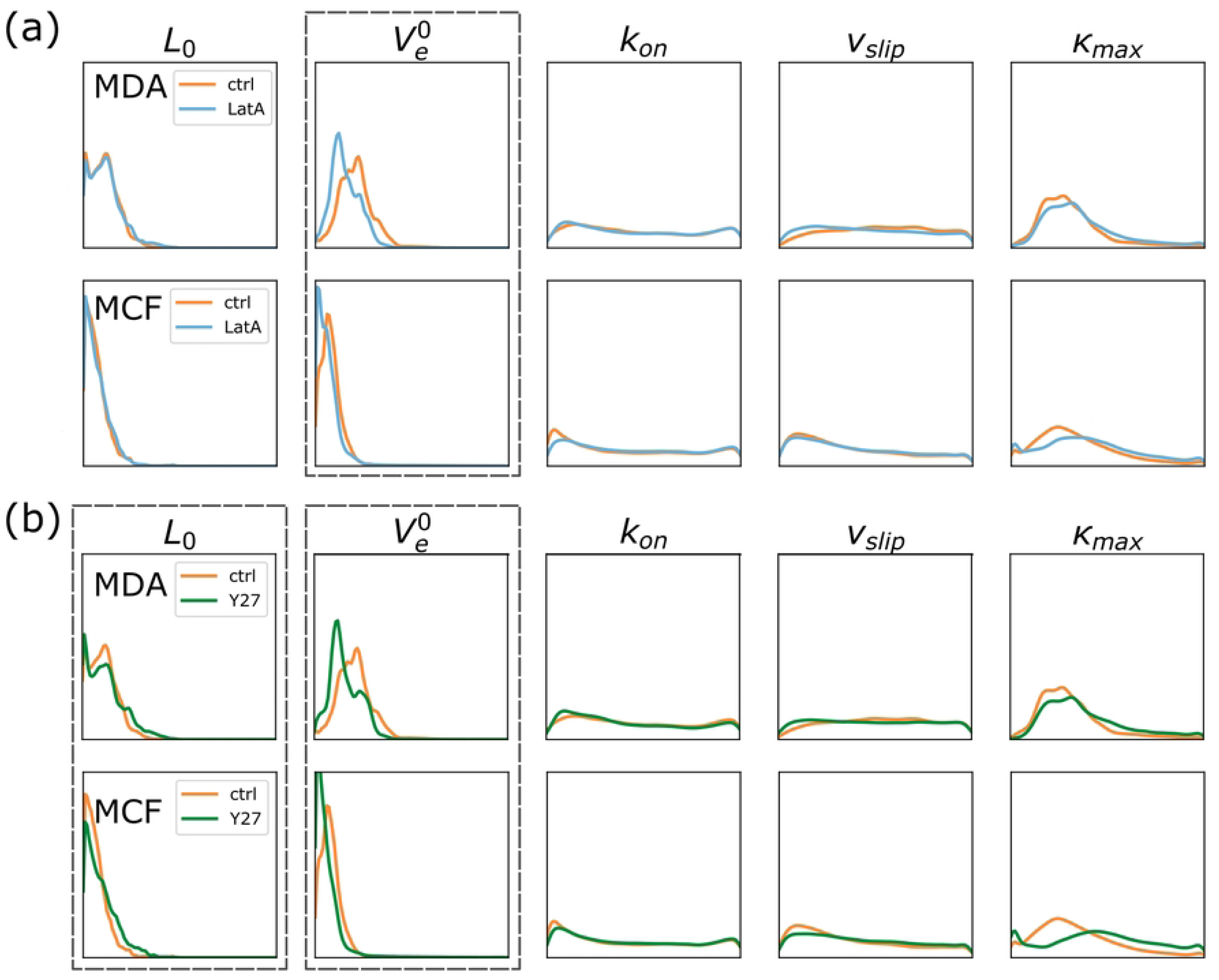
Effect of inhibitors on model parameters as inferred by SBI. Ensemble posterior distribution of model parameters of experimental cell trajectories using SBI. (a) Latrunculin A significantly decreases the actin network extension rate V_e_ ^0^ for both MDA-MB-231 and MCF-10A cells while leaving all other parameters unchanged. (b) A similar effect can be observed for Y27632 treatment. Additionally, the resting protrusion length L_0_ is shifted towards larger values. For each trajectory in a given population we sampled 1000 different points in parameter space. Here, the ensemble of all sampled points for all trajectories of a given population is shown. MDA experiments, 5 replications, N_MDA_ctrl_ = 85, N_MDA_LatA_ = 129, N_MDA_Y27_ = 96; MCF experiments, 4 replications, N_MCF_ctrl_ = 301, N_MCF_LatA_ = 465, N_MCF_Y27_ = 507

The inferred changes in the parameter probability distribution are in good agreement with the known action of the inhibitors. For Latrunculin A we expect a decrease of the actin network extension rate V_e_^0^ as Latrunculin A specifically binds to the barbed sides of the actin filaments. In our model all other parameters are independent of actin polymerisation and should not be affected by Latrunculin A. The inferred distribution functions are in excellent agreement with expectation. The Rho/ROCK pathway is an essential regulatory control element in mesenchymal cell migration with more complex consequences (40,48). ROCK phosphorylates LIM kinases that in turn phosphorylate cofilin. Cofilin is a key regulator of actin turnover that depolymerizes f-actin. By phosphorylating cofilin, ROCK/LIMK effectively inhibits actin depolymerization. Additionally, ROCK increases myosin II activity and contractility by inhibiting the dephosphorylation of myosin light chain (MLC). Furthermore, Rho and ROCK are involved in the regulation of cell-substratum adhesion via the promotion of focal-adhesion assembly and turnover (48). Srinivasan et al. observed that Y-27632 induced inhibition of ROCK in healthy primary keratinocytes (HPKs) and epidermal carcinoma cell line (A-431 cells) resulted in loss of migration, contractility, focal adhesions, and stress fibers (50). Our SBI analysis shows that Y-27632 reduces the polymerisation rate and extends the resting length of cells, most likely due to loss of contractility, and hence is in good agreement with the general understanding of Rho/ROCK signaling. It is surprising that Y-27632 does not lead to interpretable changes of the focal adhesion parameters k_on_ and κ_max_ as would be expected from the reported action of the ROCK inhibitor. However, these parameter distributions seem to be too broad and insensitive to show an effect of the treatments. It should be noted that in the case of the characterization of the two cell lines, though, the focal adhesion parameters show significant differences, Fig 6. Importantly, the fact that both cell lines react to the same treatments in a consistent fashion hints towards an underlying mechanism shared by both cell lines. In conclusion, we show that SBI specifically retrieves the effect of the inhibitors Latrunculin A and Y-27632 in an interpretable parameter space.

## Discussion

In this paper we studied the application of simulation-based inference (SBI) to estimate parameters of a mechanistic model for cell motility. We used an automated time-lapse imaging platform to collect a large number of trajectories of cells in 1D confinement for two different cell lines and different cytoskeletal inhibitors. The trajectories exhibit significant features showing defined migration states as well as meaningful rates of locomotion and oscillatory behavior. All these features are reproduced in principle by a previously published mechanistic biophysical model. The key question remaining, however, is which parameter set quantitatively captures the dynamics of observed cell trajectories in best agreement with the data. In this context we showed that SBI, once trained and calibrated, successfully infers best estimates of parameter sets of our mechanistic model. The approach is capable of characterizing migratory phenotypes in terms of parameter distributions and to assess effects of inhibitors.

We identified limitations of the approach in terms of the dimensionality of the parameter space and introduced a reduced free parameter space. In general, more parameters should be inferable, if the data set of trajectories contain sufficient information and less noise. As shown in this work, trajectories are noisy, comprising both extrinsic as well as intrinsic noise sources. Inhomogeneities in the FN lanes arguably are sources of external noise and hence cell motility on truly homogenous lanes is likely to exhibit improved parameter estimation. Moreover, we expect that expansion of the data basis by increasing both spatial and temporal resolution would further improve the SBI approach. However, in our experiments an optimal compromise of spatio-temporal resolution and number of cell trajectories was chosen. The most relevant experimental specification for data quality is the length of individual trajectories. If the trajectory is too short it does not provide the information content necessary to infer model parameters confidently. Yet, longer trajectories are limited by cell division cycle at the latest. A larger number of trajectories of the same length, however, does not necessarily improve SBI’s performance in characterizing population ensembles. In future work it will be essential to increase the dimensionality of trajectories by monitoring additional measures. Based on sensitivity analysis of the biophysical model, quantities such as the actin retrograde flow velocity or focal adhesion density would significantly increase the precision of SBI as we demonstrate with simulated data in the SI. A closer look into the summary statistics of the trajectories might elucidate which features are the most relevant for inference.

High-throughput motility assays are instrumental to extract cell specific properties. Standardized confinement, as for example in The First World Cell Race by Maiuri et al., has already been used for comparative characterization of speed and persistence for a large variety of cell lines (2,3,50–52). In contrast to model free AI based classification, SBI builds on a mechanistic model inferring interpretable features of motile cell behavior. Automated cell platforms using SBI with generally accepted mechanistic models might generate standardized parameter data bases potentially paving the way to new discoveries in cell mechanics, pharmaceutical and potentially clinical studies (55,56). Clearly the SBI approach presented here is applicable to other models related to cell motility. For example detailed models of cell protrusion dynamics exist that reproduce protrusion oscillations and traveling wave actin dynamics at shorter time scales than shown here (57,58). In principle, any complex dynamic phenomenon that can be measured with sufficient statistics and described with non-linear partial differential equations, qualifies for SBI.

Future SBI-based approaches might also be used to assess the degree of agreement of competing cell models with data in terms of posterior distribution functions. Biophysical models evolve over time, generally becoming more detailed. SBI would allow to challenge competing models and discuss more subtle additions of model components. The data driven SBI analysis of cell trajectories proposed here combines hypothesis based modeling with AI-supported analysis and hence is most appealing to the advancement of our understanding of locomotion.

## Methods

### Experimental Methods

#### Cell culture

We cultured MDA-MB-231 cells that had been stably transduced with histone-2B mCherry (gift from Timo Betz, University of Göttingen, Germany) and MCF-10A cells (obtained from ATCC, Manassas, VA, USA) in Leibovitz’s CO_2_-buffered L-15 medium with 2 mM Glutamax (Thermo Fisher Scientific, Waltham, MA, USA) at 37°C. The growth medium for MDA-MB-231 cells was supplemented by 10% fetal bovine serum (Thermo Fisher) and the medium for MCF-10A cells by 5% horse serum (Merck, Darmstadt, Germany), human epidermal growth factor (Merck), hydrocortisone (Merck), cholera toxin (Merck) and Insulin (Merck). We passaged cells every 2–3 days using Accutase (Thermo Fisher).

For experiments, we seeded about 5,000 cells per dish. After 2–3 h, cells adhered to the micropatterns and we exchanged the medium with medium containing 25nM Hoechst 33342 (invitrogen, Waltham, MA, USA) and treatment factors. The treatment factors were 0.1μM Latrunculin A (EMD millipore, Burlington, MA, USA), 30μM Y-27632 (Sigma Aldrich) and 0.3% dimethyl sulfoxide (Life Technologies, Darmstadt, Germany) as control.

#### Micropatterning

We produced the micropatterns on a Primo system (Alvéole, France) as described by Melero et al. (9). In brief, we designed micropatterns consisting of 15μm wide Fibronectin lanes with a spacing of 73μm using the vector graphics software Inkscape (inkscape.org). We conjugated human Fibronectin (yo-proteins, Sweden) with Alexa Fluor 647 NHS-ester (Thermo Fisher). We determined the concentration of the labeled protein with a NanoDrop spectrophotometer (Thermo Fisher) and confirmed the results with a Coomassie Bradford assay (Thermo Fisher). We passivated imaging dishes with polymer coverslip bottoms (ibidi, Germany) with PLL (Sigma Aldrich) and conjugated the PLL with PEG (Laysan Bio, Arab, AL, USA). Afterwards we added photoactive PLPP gel (Alvéole) and illuminated the shape of our micropatterns onto the cover slip using the UV-beam of the Primo device. Next, we washed the dishes and incubated with the labeled Fibronectin solution. Lastly, the dishes were washed once more with PBS before we seeded cells onto the patterned cover slip bottom.

#### Microscopy

We performed time-lapse imaging on an inverted fluorescence microscope (Nikon Eclipse Ti, Nikon, Tokyo, Japan) equipped with an XY-motorized stage, Perfect Focus System (Nikon), and a heating chamber (Okolab, Pozzuoli, Italy) set to 37C. We set up an acquisition protocol to sequentially scan and image fields of view using the motorized stage, the Perfect Focus System, a 10 CFI Plan Fluor DL objective (Nikon), a CMOS camera (PCO edge 4.2, Excelitas PCO, Kelheim, Germany) and the acquisition software NIS Elements (Nikon). Before the start of the time-lapse measurement, we took epifluorescence images of the FN patterns. Phase-contrast images of the cells and epifluorescence images of their nuclei were then taken for 48 h at 10 min or 2min intervals as indicated. Intervals of 10 min allowed scanning of 13×13=169 fields of view, while intervals of 10 min allowed 8×8=64 fields of view. A temporal resolution of 2min proved optimal to capture the full extent of the migration dynamics as described here while still allowing for a sufficient number of fields of view to achieve the required statistics.

#### Image analysis

We used an in-house built <u>data pipeline</u> to extract cell trajectories from raw time-lapse experimental images. The pipeline first detects the position of each fluorescent FN lane on each of the microscope’s fields of view. Next, it uses <u>cellpose</u> (19,58) to segment each individual cell and <u>trackpy</u> (61,62) to track the fluorescent nuclei. Each single nucleus trajectory is assigned a corresponding cell mask to obtain a time-lapse of the cell’s 2D shape. The information of the cell’s shape and position is combined with the position of the fibronectin lanes to calculate the rearmost and frontmost position of the cell along the corresponding FN lane. The output is a dataset with several thousand trajectories (front, back and nucleus position) of single cells for each experiment. Finally, the data are filtered to ensure a dataset that consists only of single cell trajectories of a length of 24h.

### Biophysical Modelling

#### The biomechanical model

We present a simplified version of the biophysical model published by Amiri et al. in (13), see **Fig 2**. The system is defined by the following force balance for the front (f), back (b) and center (c) of the cell:

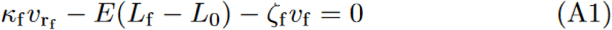

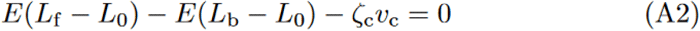

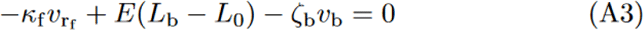

We reduced the number of parameters compared to Amiri et al. by both simplifying the model and assuming that certain parameters are fixed, see SI. To incorporate the effect of noise in the cell’s position due to both measurement and cellular factors, we added an additional source of noise to our trajectories, see next section. After these simplifications, we are left with 10 parameters and two noise amplitudes. These 10 parameters characterize a simulated cell.

We then split the remaining parameters into three possible categories: Fixed parameters which we assume to be constant for all conditions, latent parameters which we assume to vary for different simulations, but which we do not try to infer, and finally variable parameters which we vary and whose posterior p(**θ**|**x**) we approximate with NDE, see Table 1.

#### External noise

The original version of our cytoskeletal model had a single source of noise. The adhesion-dynamics κ were modeled with a Langevin equation dκ/dt = f(k)+η(t). This source of noise leads to transitions in the cell’s dynamic states. However, the shape of the simulated cell’s position **x(t)** is much smoother than experimental trajectories. The rough shape of a cell’s position in experimental trajectories can be explained by various different reasons. First, the roughness can originate from the measurement imprecisions such as the microscope’s resolution or the segmentation of cell contours (see section Image Analysis). Second, the roughness of the experimental trajectories can originate from lower-level processes that do not enter our cytoskeletal model. This dissimilarity between simulated trajectories and experimental trajectories leads to complications in simulation-based inference (SBI). The neural posterior estimator learns specific smooth features of the simulated trajectories, and performs very well in inferring simulation parameters. These smooth features are not present in simulated trajectories, so the posterior estimator cannot infer parameters of experimental trajectories. To overcome this issue, we added an additional noise source: external noise. We simply added Gaussian noise to the simulated trajectories:

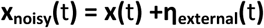

This ensured that the neural posterior estimator could not learn the smooth features in the simulated trajectories, leading to a better performance on experimental trajectories.

### Simulations

The biomechanical model presented in **Fig 2** is implemented using the Euler Method in Julia to enable fast simulations (ca. 10ms per 1 hour trajectory per CPU core) (63). The <u>source code</u> is publicly available and includes a desktop application to simulate trajectories by tweaking the values of the model’s parameters and the variables’ initial values.

### Neural Posterior Density Estimator (NDE)

We use the open source python package “sbi” developed by Tejero-Cantero et al. at the Macke lab (43) to infer the posterior distribution of model parameters of single cells given their 1D trajectories: p(**θ|x**). The algorithm implemented in the package is based on work by Greenbert et al. (35) to learn p(**θ**|**x**) directly without computing the likelihood p(**x**|**θ**). Here, the posterior is approximated by a parameterized family of functions q**_ψ_** so that p(**x**|**θ**) ≈ q**_ψ_**(**θ**). The distribution parameters **ψ** given a trajectory **x** are learned by a neural network with weights **φ**: **F** (**x**, **φ**) = **ψ**. The training of the neural network is schematically shown in **Fig 3**. We start by sampling a set of model parameters from the prior: {**θ**_j_} ∼ p(**θ**). We then simulate a trajectory for each sampled parameter set to build a dataset: {(**θ**_j_, **x**_j_)}. The neural network **F**(**x**, **φ**) learns the posterior distribution by adapting its weights **φ** to maximize the log-likelihood of true parameters given their corresponding simulated trajectories:

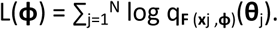

Our neural network for density estimation is composed of two main components. First, an embedding in the form of a convolutional neural network (CNN) reduces the dimensionality of our input vector, i.e. a cell trajectory, and extracts features. Then, the features obtained by the CNN are fed to a neural spline flow network (64). The input layer of the CNN was modified from being one-dimensional to being two-dimensional to better accommodate the interconnected nature of the three time serieses (front, back, nucleus) that constitute a single trajectory. This way, relations between the positions of the same cell are better preserved.

## Supporting information

### S1 Supporting information

**S1 Fig. Inference of 10 free parameters leads to loss of identifiability.** The posterior probability p(**θ**|**x**) is inferred by a neural density estimator that was learnt to infer 10 free parameters. The plots on the diagonal show the posterior distribution for each individual parameter p(θ_i_|x), while the plots in the right hand corner show the distribution for each pair of parameters p(θ_ij_|x). Vertical gray lines in the plots on the diagonal and white crosses in the plots on the off-diagonal represent the values that were used for the simulated trajectory. The posterior distributions are smeared out across the range of the prior distribution which means that the NDE can’t infer the true parameter set precisely.

**S2 Fig. Quality of the NDE** Examples of the rank statistics for 1023 simulations (N=23). The rank statistics for the N_sim_ simulations should be uniformly distributed and fall within the gray area. The parameters V_e_^0^, k_on_, v_slip_ and ⍰_max_ can be considered as being well calibrated while the posterior estimation for L_0_ is somewhat under-confident.

**S3 Fig. New variables improve SBI performance.** The posterior p(**θ**|**x**) was inferred by our trained NDPE. (a) We compare a posterior trained with the three cellular positions as input. (b) Here, the input variables contain not only the cellular positions but also the actin retrograde flows v_r,f_, v_r,b_. (c) This plot depicts a posterior trained on the cellular positions plus the adhesion dynamics κ_f_ and κ_b_. (d) Finally, the posterior if the input contains the cellular positions, the actin retrograde flows and the adhesion dynamics. The sharpening of the posterior estimator with the addition of observed variables suggests that one could characterize migrating cells much more precisely by adding further readouts to the experimental tracking.

## References

1. Matsuda T, Sugawara T. Control of cell adhesion, migration, and orientation on photochemically microprocessed surfaces. J Biomed Mater Res. 1996;32(2):165–73.

2. Maiuri P, Rupprecht JF, Wieser S, Ruprecht V, Bénichou O, Carpi N, et al. Actin flows mediate a universal coupling between cell speed and cell persistence. Cell. 2015 Apr 9;161(2):374–86.

3. Maiuri P, Terriac E, Paul-Gilloteaux P, Vignaud T, McNally K, Onuffer J, et al. The first World Cell Race. Curr Biol. 2012 Sep 11;22(17):R673–5.

4. Schreiber C, Segerer FJ, Wagner E, Roidl A, Rädler JO. Ring-Shaped Microlanes and Chemical Barriers as a Platform for Probing Single-Cell Migration. Sci Rep. 2016 May 31;6.

5. Ljepoja B, Schreiber C, Gegenfurtner FA, García-Roman J, Köhler B, Zahler S, et al. Inducible microRNA-200c decreases motility of breast cancer cells and reduces filamin A. PLOS ONE. 2019 Nov 20;14(11):e0224314.

6. Schuster SL, Segerer FJ, Gegenfurtner FA, Kick K, Schreiber C, Albert M, et al. Contractility as a global regulator of cellular morphology, velocity, and directionality in low-adhesive fibrillary micro-environments. Biomaterials. 2016 Sep 1;102:137–47.

7. Ruprecht V, Monzo P, Ravasio A, Yue Z, Makhija E, Strale PO, et al. How cells respond to environmental cues – insights from bio-functionalized substrates. Ewald A, editor. J Cell Sci. 2017 Jan 1;130(1):51–61.

8. DiMilla P, Stone J, Quinn J, Albelda S, Lauffenburger D. Maximal migration of human smooth muscle cells on fibronectin and type IV collagen occurs at an intermediate attachment strength. J Cell Biol. 1993 Aug;122(3):729–37.

9. Barnhart EL, Lee KC, Keren K, Mogilner A, Theriot JA. An Adhesion-Dependent Switch between Mechanisms That Determine Motile Cell Shape. PLoS Biol. 2011 May;9(5):e1001059.

10. Huttenlocher A, Ginsberg MH, Horwitz AF. Modulation of cell migration by integrin-mediated cytoskeletal linkages and ligand-binding affinity. J Cell Biol. 1996 Sep 15;134(6):1551–62.

11. Palecek SP, Loftus JC, Ginsberg MH, Lauffenburger DA, Horwitz AF. Integrin-ligand binding properties govern cell migration speed through cell-substratum adhesiveness. Nature. 1997 Feb;385(6616):537–40.

12. Schreiber C, Amiri B, Heyn JCJ, Rädler JO, Falcke M. On the adhesion–velocity relation and length adaptation of motile cells on stepped fibronectin lanes. Proc Natl Acad Sci. 2021 Jan 26;118(4):e2009959118.

13. Amiri B, Heyn JCJ, Schreiber C, Rädler JO, Falcke M. On multistability and constitutive relations of cell motion on fibronectin lanes. Biophys J. 2023 Feb 3;122(5):753–66.

14. Ron JE, Monzo P, Gauthier NC, Voituriez R, Gov NS. One-dimensional cell motility patterns. Phys Rev Res. 2020 Aug 11;2(3):033237.

15. Hennig K, Wang I, Moreau P, Valon L, DeBeco S, Coppey M, et al. Stick-slip dynamics of cell adhesion triggers spontaneous symmetry breaking and directional migration of mesenchymal cells on one-dimensional lines. Sci Adv. 2020;6(1):1–13.

16. Drozdowski OM, Ziebert F, Schwarz US. Optogenetic control of migration of contractile cells predicted by an active gel model. Commun Phys. 2023 Jun 30;6(1):1–12.

17. Sens P. Stick–slip model for actin-driven cell protrusions, cell polarization, and crawling. Proc Natl Acad Sci. 2020 Oct 6;117(40):24670–8.

18. Ronneberger O, Fischer P, Brox T. U-net: Convolutional networks for biomedical image segmentation. In: Medical Image Computing and Computer-Assisted Intervention (MICCAI) [Internet]. Springer; 2015. p. 234–41. Available from: http://lmb.informatik.uni-freiburg.de/Publications/2015/RFB15a

19. Stringer C, Wang T, Michaelos M, Pachitariu M. Cellpose: a generalist algorithm for cellular segmentation. Nat Methods. 2021 Jan;18(1):100–6.

20. Ma X, Dagliyan O, Hahn KM, Danuser G. Profiling cellular morphodynamics by spatiotemporal spectrum decomposition. PLOS Comput Biol. 2018 Feb 8;14(8):e1006321.

21. Tsai AG, Glass DR, Juntilla M, Hartmann FJ, Oak JS, Fernandez-Pol S, et al. Multiplexed single-cell morphometry for hematopathology diagnostics. Nat Med. 2020 Mar;26(3):408–17.

22. Masaeli M, Gupta D, O’Byrne S, Tse HTK, Gossett DR, Tseng P, et al. Multiparameter mechanical and morphometric screening of cells. Sci Rep. 2016 Dec 2;6(1):37863.

23. Hölscher DL, Bouteldja N, Joodaki M, Russo ML, Lan YC, Sadr AV, et al. Next-Generation Morphometry for pathomics-data mining in histopathology. Nat Commun. 2023 Jan 28;14(1):470.

24. Bakal C, Aach J, Church G, Perrimon N. Quantitative Morphological Signatures Define Local Signaling Networks Regulating Cell Morphology. Science. 2007 Jun 22;316(5832):1753–6.

25. Moen E, Bannon D, Kudo T, Graf W, Covert M, Van Valen D. Deep learning for cellular image analysis. Nat Methods. 2019 Dec;16(12):1233–46.

26. Vasilevich A, de Boer J. Robot-scientists will lead tomorrow’s biomaterials discovery. Curr Opin Biomed Eng. 2018 Jun 1;6:74–80.

27. Salek M, Li N, Chou HP, Saini K, Jovic A, Jacobs KB, et al. COSMOS: a platform for real-time morphology-based, label-free cell sorting using deep learning. Commun Biol. 2023 Sep 22;6(1):1–11.

28. Mavropoulos A, Johnson C, Lu V, Nieto J, Schneider EC, Saini K, et al. Artificial Intelligence-Driven Morphology-Based Enrichment of Malignant Cells from Body Fluid. Mod Pathol. 2023 Aug 1;36(8):100195.

29. Yin Z, Sadok A, Sailem H, McCarthy A, Xia X, Li F, et al. A screen for morphological complexity identifies regulators of switch-like transitions between discrete cell shapes. Nat Cell Biol. 2013 Jul;15(7):860–71.

30. Cranmer K, Brehmer J, Louppe G. The frontier of simulation-based inference. Proc Natl Acad Sci. 2020 Dec;117(48):30055–62.

31. Gonçalves PJ, Lueckmann JM, Deistler M, Nonnenmacher M, Öcal K, Bassetto G, et al. Training deep neural density estimators to identify mechanistic models of neural dynamics. Huguenard JR, O’Leary T, Goldman MS, editors. eLife. 2020 Sep 17;9:e56261.

32. Oesterle J, Behrens C, Schröder C, Hermann T, Euler T, Franke K, et al. Bayesian inference for biophysical neuron models enables stimulus optimization for retinal neuroprosthetics. Borst A, Huguenard JR, Borst A, Fairhall AL, editors. eLife. 2020 Oct 27;9:e54997.

33. Bittner SR, Palmigiano A, Piet AT, Duan CA, Brody CD, Miller KD, et al. Interrogating theoretical models of neural computation with emergent property inference. Huguenard JR, O’Leary T, Goldman MS, editors. eLife. 2021 Jul 29;10:e56265.

34. Tolley N, Rodrigues PLC, Gramfort A, Jones SR. Methods and considerations for estimating parameters in biophysically detailed neural models with simulation based inference. PLOS Comput Biol. 2024 Feb 26;20(2):e1011108.

35. Greenberg DS, Nonnenmacher M, Macke JH. Automatic Posterior Transformation for Likelihood-free Inference. In: Proceedings of the 36 th International Conference on Machine Learning. 2019.

36. Papamakarios G, Murray I. Fast \epsilon -free Inference of Simulation Models with Bayesian Conditional Density Estimation. In: Advances in Neural Information Processing Systems [Internet]. Curran Associates, Inc.; 2016 [cited 2024 Apr 2]. Available from: https://proceedings.neurips.cc/paper_files/paper/2016/hash/6aca97005c68f1206823815f66102863-Abstract.html

37. Lueckmann JM, Goncalves PJ, Bassetto G, Öcal K, Nonnenmacher M, Macke JH. Flexible statistical inference for mechanistic models of neural dynamics. In: Advances in Neural Information Processing Systems [Internet]. Curran Associates, Inc.; 2017 [cited 2024 Apr 2]. Available from: https://proceedings.neurips.cc/paper/2017/hash/addfa9b7e234254d26e9c7f2af1005cb-Abstract.html

38. Deistler M, Goncalves PJ, Macke JH. Truncated proposals for scalable and hassle-free simulation-based inference. Adv Neural Inf Process Syst. 2022 Dec 6;35:23135–49.

39. Coué M, Brenner SL, Spector, Ilan (National Heart, Lung, and Blood Institute M, Korn D. Inhibition of actin polymerization by latrunculin A. FEBS Lett. 1987;213(2):316–8.

40. Bolado-Carrancio A, Rukhlenko OS, Nikonova E, Tsyganov MA, Wheeler A, Garcia-Munoz A, et al. Periodic propagating waves coordinate RhoGTPase network dynamics at the leading and trailing edges during cell migration. Mogilner A, Walczak AM, Edelstein-Keshet L, editors. eLife. 2020 Jul;9:e58165.

41. Gardel ML, Sabass B, Ji L, Danuser G, Schwarz US, Waterman CM. Traction stress in focal adhesions correlates biphasically with actin retrograde flow speed. J Cell Biol. 2008 Dec 15;183(6):999–1005.

42. Craig EM, Stricker J, Gardel M, Mogilner A. Model for adhesion clutch explains biphasic relationship between actin flow and traction at the cell leading edge. Phys Biol. 2015 May;12(3):035002.

43. Tejero-Cantero A, Boelts J, Deistler M, Lueckmann JM, Durkan C, Gonçalves PJ, et al. sbi: A toolkit for simulation-based inference. J Open Source Softw. 2020 Aug 21;5(52):2505.

44. Kashman Y, Groweiss A, Shmueli U. Latrunculin, a new 2-thiazolidinone macrolide from the marine sponge latrunculia magnifica. Tetrahedron Lett. 1980;21(37):3629–32.

45. Spector, Ilan (National Heart, Lung, and Blood Institute M, Shochet, Nava R (National Heart, Lung, and Blood Institute M, Kashman Y (Tel AU, Groweiss A (Tel AU. Latrunculins: novel marine toxins that disrupt microfilament organization in cultured cells. Science. 1983 Feb 4;219(4584):493–5.

46. Kuwahara K, Saito Y, Nakagawa O, Kishimoto I, Harada M, Ogawa E, et al. The effects of the selective ROCK inhibitor, Y27632, on ET-1-induced hypertrophic response in neonatal rat cardiac myocytes – possible involvement of Rho/ROCK pathway in cardiac muscle cell hypertrophy. FEBS Lett. 1999;452(3):314–8.

47. Uehata M, Ishizaki T, Satoh H, Ono T, Kawahara T, Morishita T, et al. Calcium sensitization of smooth muscle mediated by a Rho-associated protein kinase in hypertension. Nature. 1997 Oct;389(6654):990–4.

48. Riento K, Ridley AJ. ROCKs: multifunctional kinases in cell behaviour. Nat Rev Mol Cell Biol. 2003 Jun;4(6):446–56.

49. Srinivasan S, Das S, Surve V, Srivastava A, Kumar S, Jain N, et al. Blockade of ROCK inhibits migration of human primary keratinocytes and malignant epithelial skin cells by regulating actomyosin contractility. Sci Rep. 2019 Dec 27;9(1):19930.

50. Renkawitz J, Kopf A, Stopp J, de Vries I, Driscoll MK, Merrin J, et al. Nuclear positioning facilitates amoeboid migration along the path of least resistance. Nature. 2019 Apr;568(7753):546– 50.

51. Kopf A, Renkawitz J, Hauschild R, Girkontaite I, Tedford K, Merrin J, et al. Microtubules control cellular shape and coherence in amoeboid migrating cells. J Cell Biol. 2020 May 7;219(6):e201907154.

52. Brückner DB, Broedersz CP. Learning dynamical models of single and collective cell migration: a review. Rep Prog Phys. 2024 Apr;87(5):056601.

53. Masuzzo P, Van Troys M, Ampe C, Martens L. Taking Aim at Moving Targets in Computational Cell Migration. Trends Cell Biol. 2016 Feb 1;26(2):88–110.

54. Wilkinson MD, Dumontier M, Aalbersberg IjJ, Appleton G, Axton M, Baak A, et al. The FAIR Guiding Principles for scientific data management and stewardship. Sci Data. 2016 Mar 15;3(1):160018.

55. Enculescu M, Sabouri-Ghomi M, Danuser G, Falcke M. Modeling of Protrusion Phenotypes Driven by the Actin-Membrane Interaction. Biophys J. 2010;98(8):1571–81.

56. Dolati S, Kage F, Mueller J, Müsken M, Kirchner M, Dittmar G, et al. On the relation between filament density, force generation, and protrusion rate in mesenchymal cell motility. Mol Biol Cell. 2018 Nov;29(22):2674–86.

57. Melero C, Kolmogorova A, Atherton P, Derby B, Reid A, Jansen K, et al. Light-Induced Molecular Adsorption of Proteins Using the PRIMO System for Micro-Patterning to Study Cell Responses to Extracellular Matrix Proteins. JoVE J Vis Exp. 2019 Oct 11;(152):e60092.

58. Pachitariu M, Stringer C. Cellpose 2.0: how to train your own model. Nat Methods. 2022 Dec;19(12):1634–41.

59. Crocker JC, Grier DG. Methods of Digital Video Microscopy for Colloidal Studies. J Colloid Interface Sci. 1996 Apr 15;179(1):298–310.

60. Allan DB, Caswell T, Keim NC, van der Wel CM, Verweij RW. soft-matter/trackpy: v0.6.1 [Internet]. Zenodo; 2023 [cited 2023 Aug 1]. Available from: https://zenodo.org/record/7670439

61. Bezanson J, Karpinski S, Shah VB, Edelman A. Julia: A Fast Dynamic Language for Technical Computing [Internet]. arXiv; 2012 [cited 2023 Aug 1]. Available from: http://arxiv.org/abs/1209.5145

62. Durkan C, Bekasov A, Murray I, Papamakarios G. Neural Spline Flows [Internet]. arXiv; 2019 [cited 2023 Aug 24]. Available from: http://arxiv.org/abs/1906.04032

63. Talts S, Betancourt M, Simpson D, Vehtari A, Gelman A. Validating Bayesian Inference Algorithms with Simulation-Based Calibration [Internet]. arXiv; 2020 [cited 2024 May 2]. Available from: http://arxiv.org/abs/1804.06788

